# Neural organoids protect engineered heart tissues from glucolipotoxicity by transferring versican in a co-culture system

**DOI:** 10.1101/2025.05.07.652597

**Authors:** Baochen Bai, Jiting Li, Ze Wang, Yuhan Yang, Jieqing He, Gonglie Chen, Yufan Zhang, Yan Qi, Zhongjun Wan, Lin Cai, Run Wang, Kai Wang, Dongyu Zhao, Jingzhong Zhang, Weihua Huang, Ronald X Xu, Mingzhai Sun, Xiao Han, Yan Liu, Donghui Zhang, Wanying Zhu, Jian Liu, Yuxuan Guo

## Abstract

Metabolic disorders could cause dysregulated glucose and lipid at the systemic level, but how inter-tissue/organ communications contribute to glucolipotoxicity is difficult to dissect in animal models. To solve this problem, myocardium and nerve tissues were modeled by 3D engineered heart tissues (EHTs) and neural organoids (NOs), which were co-cultured in a generalized medium with normal or elevated glucose/fatty acid contents. Morphology, gene expression, cell death and functional assessments detected no apparent alterations of EHTs and NOs in co-culture under normal conditions. By contrast, NOs significantly ameliorated glucolipotoxicity in EHTs. Transcriptomic and protein secretion assays identified the extracellular matrix protein versican as a key molecule that was transferred from NOs into EHTs in the high-glucose/fatty acid condition. Recombinant versican protein treatment was sufficient to reduce glucolipotoxicity in EHTs. Adeno-associated virus-delivered versican overexpression was sufficient to ameliorate cardiac dysfunction in a murine model of diabetic cardiomyopathy. These data provide the proof-of-concept evidence that inter-tissue/organ communications exist in the co-culture of engineered tissues and organoids, which could be systemically studied to explore potential pathological mechanisms and therapeutic strategies for multi-organ diseases in vitro.

## 1. Introduction

The dysregulation of glucose and fatty acids is commonly observed in metabolic diseases such as obesity and diabetes. High glucose and fatty acid levels could cause compound pathological insults termed glucolipotoxicity, which was initially referred to their adverse impacts on pancreatic β-cells^1^. With a more systemic view of metabolic syndrome, glucolipotoxicity is believed to occur in other organs particularly in the cardiovascular system^2–4^ and the nervous system^5–7^. For example, diabetic cardiomyopathy and diabetic neuropathy are considered as major complications in diabetes^8^. Despite intensive investigations of these comorbidities in individual organs, a systemic understanding of inter-tissue/organ communications is lacking due to the absence of a simplified model system.

Tissue engineering and organoid technologies are recently emerging as powerful tools to model human diseases^9^. On one hand, they assemble multiple cell types in 3D to mimic key structures and functions of a give tissue or organ, better representing pathophysiological conditions than canonical monolayer cell culture. On the other hand, they are more simplified than animal models, directing the research focus on selected tissues or organs while avoiding interference by other organs in the animals. With the recent surge of co-culture^10–14^ followed by assembloid^15, 16^ studies, engineered tissues and organoids hold the potential to build microphysiological systems (MPSs) in vitro for advanced pathological and pharmaceutical studies at the system level.

An essential prerequisite in the effort to build and apply MPSs is to understand the biological effects about the coexistence of two or more engineered tissues and organoids in the same dish. To date, such studies have been limited to anatomically or developmentally related organoids, such as the interactions between organoids from different brain subdomains^15–17^ or distinct heart chambers^18^. The culture conditions to maintain organoids of more distal origins in a dish remain to be established. Whether inter-tissue/organoid communications contribute to disease phenotypes at the system level and harbor pathologically meaningful mechanisms are also unknown.

To answer these questions, this study established a co-culture system between neural organoids (NOs) and engineered heart tissues (EHTs). Exploration of the impacts of NOs on EHTs in both normal and glucolipotoxicity conditions revealed inter-organoid communications that could potentially uncover novel therapeutic strategies.

## 2. Materials and Methods

### 2.1 Rat primary cardiac cells

Hearts were extracted from 1-day-old neonatal Sprague-Dawley rats, and washed in Hank’s Balanced Salt Solution (HBSS) (PB190321, Procell), and then enzymatically digested with 5 mL lysis buffer containing 0.1% trypsin (T8150, Solarbio) and 0.056% collagenase type II (LS004177, Worthington Biochemical) for 4 minutes at 37°C. The above steps were repeated 10 to 13 times until the hearts were completely dissociated.

Cells collected after each digestion were centrifuged at 300g for 5 min, and resuspended in pre-plate media consisting of high glucose Dulbecco’s Modified Eagle’s Medium (DMEM) (HK2109.07, HUANKE), 10% fetal bovine serum (FBS) (SE100-011, VISTECH) and 1% penicillin-streptomycin solution (BC-CE-007, Bio-channel). After then, cell suspension was passed through a 40 μm cell strainer (93040, SPL life sciences) and preplated on a cell culture dish for 2 hours to remove the most of fibroblasts and enrich the fraction of cardiomyocytes. The suspended cells were collected to determine the proportion of troponin T2 (TNNT2) positive cardiomyocytes by flow cytometry, and for EHT construction or monolayer culture.

For Ca^2+^ transient measurements, neonatal rat cardiomyocytes were incubated with 5 μM Fluo-4 AM (BioTss) diluted in medium at 37 °Cfor 60 min. Ca^2+^ transient videos were acquired using a Leica THUNDER Imager DMi8 at a speed of 10 frames per second by a 20 × objective. An Okolab Cage Incubation system maintained the temperature at 37 °C during imaging. Ca^2+^ transient amplitudes and beat rates were analyzed using ImageJ and MATLAB.

### 2.2 hESC culture and differentiation into cardiac cells

Stem cell research was performed according to the general requirements for stem cells^19^ and embryonic stem cells^20^. H7 human embryonic stem cells^21^ (WiCell, WAe007-A, NIHhESC-10-0061) were seeded in 6-well plates which are precoated with matrigel solution (CA3003100, Cellapy), and cultured in PGM1 medium (CA1007500, Cellapy) at 37°C in humidified air with 5% CO2. Cells were passaged with digestion solution (CA1023100, Cellapy) at a 1: 6 ratio when 80%-100% confluence was achieved.

Human Cardiomyocyte Differentiation Kit (CA2004500, Cellapy) were used for cardiac cells differentiation as previously described^22^. To be specific, hESCs were cultured in RPMI 1640 medium supplemented with B27 when 80%-95% confluence was achieved on day 0. Then, 5 µM CHIR99021 was added to the culture medium in the first two days. Subsequently, 5 µM IWR-1-endo was added to the culture medium from day 2 to day 4. Ten days later, the medium was replaced with cardiomyocyte maintenance medium (CA2015002, Cellapy). The fresh medium was replaced every two days. On day 30, differentiated cardiac cells were analyzed by flow cytometry to determine the proportion of TNNT2 positive cardiomyocytes. Meanwhile, differentiated cardiac cells at day 30 were digested with Human Cardiomyocyte Digestion Kit (CA2004500, Cellapy) to form hEHT.

### 2.3 EHT construction and force analysis

The EHT was constructed as previously described^23–25^. Polydimethylsiloxane (PDMS) solution was prepared by blending PDMS base with PDMS curing agent (DC184, Dow Corning) in a volumetric ratio of 10:1 and degassed for 1 hour in a vacuum desiccator.

PDMS solution was poured into a positive mold, degassed in a vacuum desiccator to remove air bubbles and cured in an oven at 56°C for at least 4 hours. The PDMS mold (negative mold) was then peeled off the positive molds and then utilized for subsequent experiments.

Before cell seeding, PDMS molds were sonicated, autoclaved and coated with 0.2% (w/v) Pluronic® F-127 (P2443, Sigma) over 2 hours. Laser-cut nylon frames (outer dimensions 14.6 mm × 11.7 mm, inner dimensions 11.7 mm × 7.8 mm) were in the PDMS mold to provide anchorage for EHTs. Next 1.2 million rat primary cardiac cells or hESC-differentiated cardiac cells and 1 U/mL thrombin (T6884, Sigma) were mixed with a fibrin-based hydrogel containing 2 mg/mL fibrinogen (F3879, Sigma) and 10% v/v matrigel (354277, Corning). This mixture was injected into the PDMS molds, and incubated at 37°C for 30 mins to allow hydrogel polymerization and attachment.

EHTs were cultured in a static incubator on the first day, and then cultured in the RPMI 1640 medium containing 0.1 mM non-essential amino acids (NEAA) (PB180424, Pricella), 0.4 mg/mL L-ascorbic acid (A8960, Sigma) and 2 mg/mL 6-aminocaproic acid (A2504, Sigma) on a shaker in the incubator. 10 µM Y27 (S104928, Selleckchem) and 100 µM BrdU (BB4261, Bestbio) were added on day 0 to inhibit the proliferation of non-myocytes. Culture medium was changed every other day. The key reagents for EHT construction are shown in Supplementary Table 1. A quality EHT with spontaneous beating is shown in Supplementary Video 1.

To assess the contractile function of EHTs, we used a custom-engineered force measurement apparatus containing a high-resolution force transducer and a computer-driven linear actuator to quantify the contractile force upon electrical field stimulation. EHTs were submerged in DMEM medium at 37°C with one end anchored to the stage and the opposing end affixed to a force transducer. Subsequently, EHTs were paced at 1.5 Hz-3 Hz for 30s of data acquisition. Strains on the EHTs were set by the actuator, and the contractile force and tension amplitudes were recorded when stretching the tissue from 0% (resting length) to 8% with 2% increments. The force data were recorded by LabScribe and analyzed by a custom MATLAB script.

### 2.4. NOs construction

Neural organoids (NOs) were generated using a well-established protocol, as reported in previous studies^26, 27^. The key reagents for NO construction are shown in Supplementary Table 1. Firstly, independent hESCs colonies after 5-7 days of culture were digested with Dispase II (D6430, Solarbio) and cultured in serum-free neural induction medium (NIM) to form embryoid bodies (EBs). Subsequently, EBs were cultured in NIM supplemented with 2 μM SB431542 (TGF-β/Smad inhibitors) (T1726, Topscience) and 2 μM dorsomorphin homolog 1 (DMH-1, BMP receptor inhibitor) (T1942, Topscience) for 7 days to induce neural differentiation. NIM medium consisted of DMEM/F12 medium (PM150313, Pricella) supplemented with 1×N2 supplement (17502001, Gibco) and 1×NEAA (PB180424, Pricella). On day 7, EBs were evenly distributed into 6-well plates and briefly cultured in NIM medium containing 10% FBS (SE100-011, VISTECH) for 10h. The medium was subsequently changed to NIM medium and then half-changed every other day. The rosette structures could be observed from day 7 to day 16. On day 16, colonies containing rosette structures were gently blown off and suspension cultured in NIM medium supplemented with 1×B27 supplement (12587001, Gibco) that was withdrawn one day later. Thereafter, the medium was half-changed every other day until day 30 for subsequent experiments.

### 2.5. EHT-NO co-culture and HGHF treatment

The EHT-NO co-culture system was constructed by transferring 2-3 NOs into medium with EHTs in 12-well plates, which were shaken on the incubator shakers. Single EHTs and EHT-NO co-culture groups were cultured in general medium containing 33 mM D-glucose (G8769, Sigma) and 200 µM palmitic acid (PA) (P5585, Sigma) for 48h to induce glucolipotoxicity as HGHF treatment. The normal glucose and normal fatty acid (NGNF) groups were cultured in general medium containing 17.4 mM D-glucose and no PA.

Preparation of HGHF medium: 45% D-glucose solution was diluted into 192.5 mM solution with ultrapure water and stored at 4°C. PA powder was dissolved in absolute ethanol to create a concentrated stock solution of 250 mM. This solution was further diluted into a 2 mM PA solution with 10% Bovine Serum Albumin (BSA) (A9418, Sigma) and stored at -20°C. The stock solutions were diluted into the general medium to prepare the final HFHG medium with desired fatty acids (200 μM) and glucose (33 mM).

Preparation of NGNF control medium: D-mannitol powder (IM0040, Solarbio) was dissolved in ultrapure water to prepare 192.5 mM concentrated solution as isotonic control for glucose. The control solution for PA was the 10% BSA solution containing equal volume absolute ethanol without PA. These concentrated solutions were dissolved into the general medium to make the final NGNF medium.

### 2.6 Organoid sectioning and immunofluorescence

Organoids were washed with PBS prior to fixation. They were then fixed for 30 minutes using 4% Paraformaldehyde (PFA), followed by gradient dehydration in the 20% and 30% sucrose solution at 4°C overnight before being embedded in Optimal Cutting Temperature (OCT). Organoids fixed in OCT-containing freezing molds were rapidly frozen and stored at temperatures below -80°C until sectioning. Leica CM3050 S Cryostat was set to a section thickness of 7 μm.

For immunostaining, sections were washed three times with PBS to remove OCT compound and blocked and permeabilized in 4% BSA supplemented with 0.1% Triton X-100 for 1 hour. Sections were incubated with 200 µL/slice of primary antibodies diluted in blocking buffer overnight at 4°C. Then the primary antibodies were washed three times with PBS, sections were incubated with fluorescent secondary antibodies for 2 hours at room temperature. After rinsing with PBS, the sections were mounted using a fluorescence mounting medium. Antibody information is listed in Supplementary Table 2. TUNEL BrightRed Apoptosis Detection Kit (A113, Vazyme Biotech) was used to detect apoptosis according to the manufacturer’s instructions. Images were taken by the Olympus FV3000 confocal microscope and quantified with ImageJ software.

### 2.7 RT-qPCR analysis

Total RNA was isolated using the TransZol Up Plus RNA Kit (ER501-01-V2, TransGene) according to the manufacturer protocols, ensuring the concurrent elimination of genomic DNA. RNA was reverse-transcribed using HiScript III All-in-one RT SuperMix (R333-01, Vazyme Biotech). RT-qPCR was performed with the AriaMx Real-Time PCR System (Agilent Technologies) and 2×Taq Pro Universal SYBR qPCR Master Mix (Q712-02, Vazyme Biotech). Relative changes on gene expression levels were normalized to *GAPDH* gene expression levels. The primer sequences are shown in Supplementary Table 3.

### 2.8 Smart-seq analysis

RNAprep Pure Micro kit (DP420, Tiangen) was used to extract total RNA from single hEHTs or NOs. Smart-Seq was next performed at the Beijing Geekgene Company with a published protocol^28^. The RNA was mixed with 0.1 μL RNase inhibitor (Clontech), 1.9 μL Triton X-100 solution (1%), 1 μL dNTP mix (10 mM), and 1 μL oligo-dT primer (5 μM). Reverse transcription was performed by further adding 0.5 μL SuperScript II reverse transcriptase (200 U/μL, Invitrogen), 0.25 μL RNase inhibitor (40 U/μL, Clontech), 2 μL Superscript II First-Strand Buffer (5×, Invitrogen), 0.5 μL DTT (0.1 M, Invitrogen), 2 μL Betain (5 M, Sigma), 0.06 μL MgCl2 (1 M, Sigma), and 0.1 μL template-switching oligos (100 μM). Reverse transcription was carried out at 25°C for 5 min, 42°C for 60 min, followed by 50°C for 30 min and 72°C for 10 min.

PCR preamplification was performed using KAPA HiFi HotStart Ready MIX (KAPA Biosystems) with 22 cycles of PCR and the IS PCR primer reduced to 50 nM (4 cycles at 98°C for 20 s, 65°C for 30 s, and 72°C for 5 min, followed by 18 cycles at 98°C for 20 s, 67°C for 15 s, and 72°C for 5 min, with a final cycle at 72°C for 5 min). Subsequently, Amplified samples were purified twice with 0.8X Ampure XP beads (Beckman, A63882). Then the construction of library was performed using KAPA Hyper Prep Kits (KK8504, Roche). Subsequently, the library was sequenced on an Illumina Novaseq 6000 platform to produce 150 bp paired-end reads.

During data analysis, FastQC software^29^ was used for quality control, and Trimmomatic (v.0.39)^30^ was used to remove low-quality reads and adapter sequences from the raw reads to obtain clean reads. The clean reads were then aligned to the human reference genome (hg38) using STAR (v.2.7.5c)^31^. Subsequently, gene expression was quantified using featureCounts (v.1.6.5)^32^ to obtain counts data, which were further subjected to bioinformatics analysis using Rstudio software.

DESeq2 package^33^ was used to analyze differentially enriched genes, and Padj < 0.05 was used as the cut-off value to identify differentially expressed genes. Principal Component Analysis (PCA) plot and MA plot were used to visualize the data distribution using Rstudio software. Gene Set Enrichment Analysis (GSEA) software^34^ was used to perform Gene Ontology (GO) analysis and Kyoto Encyclopedia of Genes and Genomes (KEGG) pathway enrichment analysis. Terms with P-value < 0.05 were considered as significant. The expression of genes included in the selected terms were normalized to Z-scores and plotted as heatmaps using GraphPad Prism 8.0.2.

### 2.9 Enzyme-linked immunosorbent assay (ELISA) for versican

Human versican ELISA kit (JL52482, Jianglai Biotech) was used to detect the concentration of versican in culture media or organoid lysates according to the manufacturer’s instructions. The Origin 2022b software was used to produce the standard curve and analyze the data.

### 2.10 Animal

Adult C57BL/6J male mice were purchased and bred in the Department of Laboratory Animal Science of Peking University Health Science Center with SPF standard. All mice were given free water and food at an ambient temperature of 20-26℃ and a light cycle of 12 h light/12 h darkness. All animal experimental procedures were approved by the Institutional Animal Care and Use Committee of Peking University (approval number DLASBD0203).

4-week-old C57BL/6J mice were randomly divided into control and DbCM group after 1 week of acclimatization. The control group was given normal control diet (NCD). Mice in DbCM group were fed with a diet with 60 kcal% fat (D12492, Research Diets) for 8 weeks, which was followed by an intraperitoneal injection of streptozotocin (STZ) at a dose of 30 mg/kg/day for 5 consecutive days. After 1 week, the tail tip blood of mice was collected to validate the fasting blood glucose level, and then the high-fat feeding was continued until the 28th week of age before the animals were used for studies.

### 2.11 Plasmids and AAV

The coding sequence of murine *Vcan* transcript variant 3 (*Vcan*-V3) was acquired at NCBI (NM_001134474) and synthesized by Tsingke Biotech, China. The AAV-CMV-GFP-LA plasmid (Addgene 206197)^35^ was used as the starting vector. Subsequently, the GFP-LA sequence was replaced by the *Vcan*-V3-HA coding sequence to generate the AAV-CMV-*Vcan*-V3-HA plasmid. AAV9 production was performed at PackGene Biotech. AAV9 was prepared as previously described^36^. AAV (1 × 10^13^ vg/kg) was intravenously injected into 28-week old mice. Four weeks later, the mice were subjected to analysis.

### 2.12 Echocardiography

Mice were anesthetized using 1-1.5% isoflurane, and echocardiography was performed with a 30 MHz MX400 transducer (Vevo 3100 ultrasound imaging system, Fujifilm VisualSonics). Left ventricular ejection fraction (EF) and fractional shortening (FS) were calculated from the parasternal short-axis sections based on M-mode images. The ratio of peak early diastolic flow filling to peak atrial systolic flow (E/A) was calculated based on Pulsed-Wave Doppler mode images. Peak early diastolic mitral annulus velocity (E′) was obtained by Tissue Doppler mode.

### 2.13 Mouse metabolic tests

For glucose tolerance test, mice were fasted for 16 h and then intraperitoneally injected with 20% glucose solution at a dose of 1 g/kg. Blood glucose levels were measured by a glucometer at 0 min, 30 min, 60 min and 90 min after injection.

For insulin tolerance test, after fasting for 4 h, mice were intraperitoneally injected with insulin solution at a dose of 0.5 U/kg. The tail vein blood was collected to detect glucose at 0 min, 30 min, 60 min and 90 min after injection.

Blood glucose was measured using Accu-Chek Performa blood glucometer. Total free fatty acid was measured using an Amplex Red Free Fatty Acid Assay Kit (Beyotime, S0215S). Total cholesterol (TC) concentration in serum of mice were detected by a TC content kit (JLT1370, Jonlnbio).

### 2.14 Western blotting

Tissue samples were washed twice with PBS and lysed in RIPA buffer (P0013B, Beyotime) containing 1% protease inhibitors (20124ES03, Yeasen). Tissue lysates were centrifuged at 12000 rpm for 30 min at 4℃. Then the supernatants were isolated and protein concentration was determined using the BCA Protein Assay kit (PC0020, Solarbio). Protein lysates were mixed with 4 × SDS-PAGE Loading Buffer and incubated at 95℃ for 5 min. 20 μg proteins were separated in a 4%-12% SDS-PAGE gel and then transferred to PVDF membrane, which was subsequently blocked with 5% milk for 1 h at room temperature. Then the membranes were incubated with primary antibodies overnight at 4°C. After washing, the membranes were incubated with secondary antibodies for 1 h at room temperature. Chemiluminescence signals were detected using a Bio-Rad imaging system.

### 2.15 Data plotting and statistical analysis

All data were presented as mean ± SEM. GraphPad Prism 8.0.2 was used for statistical analysis. Unpaired t test was used for comparison between two groups. More than two groups were subjected to one-way ANOVA with Tukey’s post hoc test. Contractile force data were analyzed by two-way ANOVA with Tukey’s post hoc test. P value of less than 0.05 was considered statistically significant.

## 3. Results

### 3.1 Establishment of an EHT-based myocardial glucolipotoxicity model

Engineered heart tissues (EHTs) were first constructed using neonatal rat primary cardiac cells (Figure 1A and Supplementary Video 1). Flow cytometry measured the fraction of TNNT2^+^ cardiomyocytes to be approximately 76.4% (Figure 1B). EHTs were subjected to cryo-sectioning followed by TNNT2 immunostaining, which validated elongated cardiomyocyte and myofibril structures (Figure 1C).

**Figure 1.**
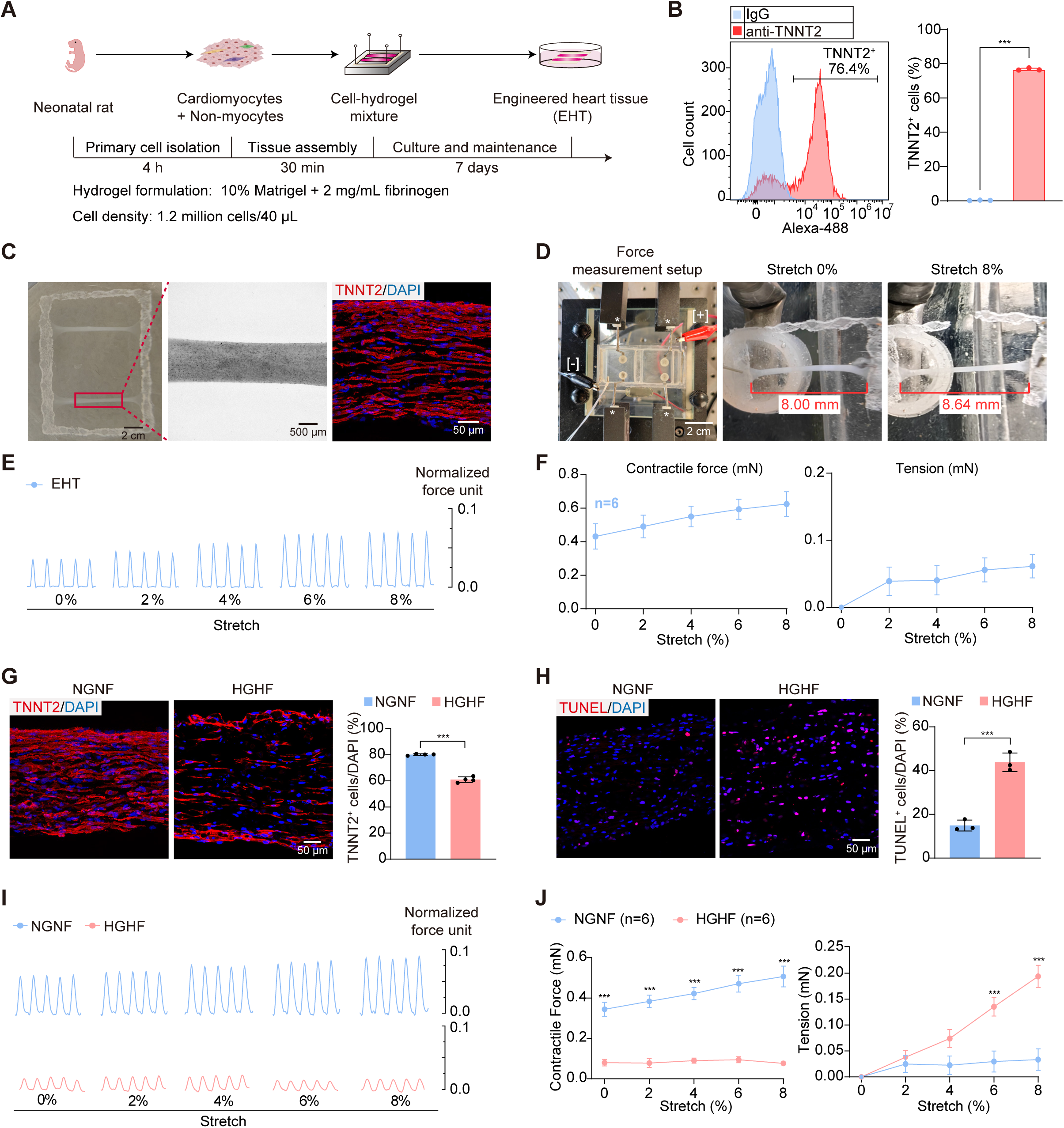
Myocardial glucolipotoxicity on engineered heart tissues (EHTs). A) A flow chart of rat EHT construction. Primary cardiac cells were extracted from neonatal rats and mixed with hydrogel, which was polymerized on a PDMS mold and attached to a rectangular Nylon frame. After 30 min solidification, EHTs were detached from the mold and cultured in medium for 7 days before experiments. B) Flow cytometry analysis of cardiomyocyte and non-cardiomyocyte fractions in primary cells. TNNT2, cardiac troponin T2. C) Representative images of EHTs. A bright-field image of two EHTs attached to a frame (left). A magnified bright-field image of a single EHT (middle). An immunofluorescence image of cardiomyocyte marker TNNT2 (red) and DAPI (blue) on a longitudinal EHT section (right). D) The image of the EHT force measurement stage (left). Electrodes for field stimulation are marked with [+] and [-]. Stars point to force sensors. The bottom of the stage was motorized to make displacement relative to the force sensor and generate stretches on EHTs. Representative unstretched (middle) and 8% stretched (right) EHTs are shown. E) Representative force traces of EHTs under 3 Hz electrical stimulation at the indicated progressive stretches. F) Quantification of the contractile force and passive tension of EHTs. G-H) Representative images and quantification of TNNT2 (G) or TUNEL staining (H) under normal glucose normal fatty acid (NGNF, 5.5 mM glucose + 0 μM palmitic acid) or high glucose high fatty acid (HGHF, 33 mM glucose + 200 μM palmitic acid) treatment for 24 h. I) Representative force traces of EHT under NGNF or HGHF treatment for 24 h. J) Quantification of the contractile force and passive tension under NGNF or HGHF treatment. Data are plotted as mean ± SEM and statistically analyzed by unpaired t test (B, G, H) or two-way ANOVA with Tukey’s post hoc test (J). ***P < 0.001.

Force measurements of EHTs were conducted using a custom platform with a motorized culture batch, force sensors and a field stimulator (Figure 1D). The two ends of EHTs were anchored to the culture batch and the force sensors, respectively, to enable measurement of contractile forces that were induced by electrical stimulation (Figure 1D). The motorized stage was moved to apply up to 8% strains on the EHTs, which resulted in increased diastolic tension and enhanced contractile force between systole and diastole (Figure 1E-F). Thus, this setup allows assessment of the frank-starling mechanism^37^, a key functional parameter of myocardium that is otherwise difficult to measure with spheric cardiac organoids^38^.

To establish a glucolipotoxicity model, 33 mM glucose and 200 μM palmitic acid were applied to EHTs (high glucose high fatty acid, HGHF). As a control, the original EHT medium (normal glucose normal fatty acid, NGNF) group involved 5.5 mM glucose and no palmitic acid. After 24h treatment, cardiomyocyte density in the HGHF group was reduced by about 20% and myofibril alignment was disrupted (Figure 1G). Terminal deoxynucleotidyl transferase dUTP nick end labeling (TUNEL) detected significantly increased apoptotic cells (Figure 1H). HGHF treatment also dramatically impaired the contractile forces while increasing diastolic tension in EHTs (Figure 1I-J).

RNA-Seq analysis of rat EHTs detected over 10,000 genes that were dysregulated by glucolipotoxicity (Supplementary Data 1 and Supplementary Figure 1A-B). Gene set enrichment analysis (GSEA) revealed increased inflammatory, cell adhesion and apoptotic processes and decreased cardiac development, cardiac contraction and oxidative respiration gene ontology (GO) terms as expected (Supplementary Figure 1C). These data support the successful establishment of an EHT-based myocardial glucolipotoxicity model.

### 3.2 Establishment of a NO-based neural glycolipotoxicity model

NOs were generated from human embryonic stem cells (hESCs) using a well-established protocol as described in previous studies^26^ (Figure 2A). As compared to hESCs, NOs at day 30 lost the expression of stem cell markers *SOX2* and *OCT4* and acquired neural markers *FOXG1* and *MAP2* (Figure 2B). RNA-Seq of single NOs by Smart-seq2^39^ detected neuron projection development, neuron migration and others as top gene ontology-biological processes (GO-BP) among the top-500 highly expressed genes (Figure 2C). The characteristic rosette structures (Figure 2D) and neural marker expression (Figure 2E) were validated by bright field and immunofluorescence imaging, respectively.

**Figure 2.**
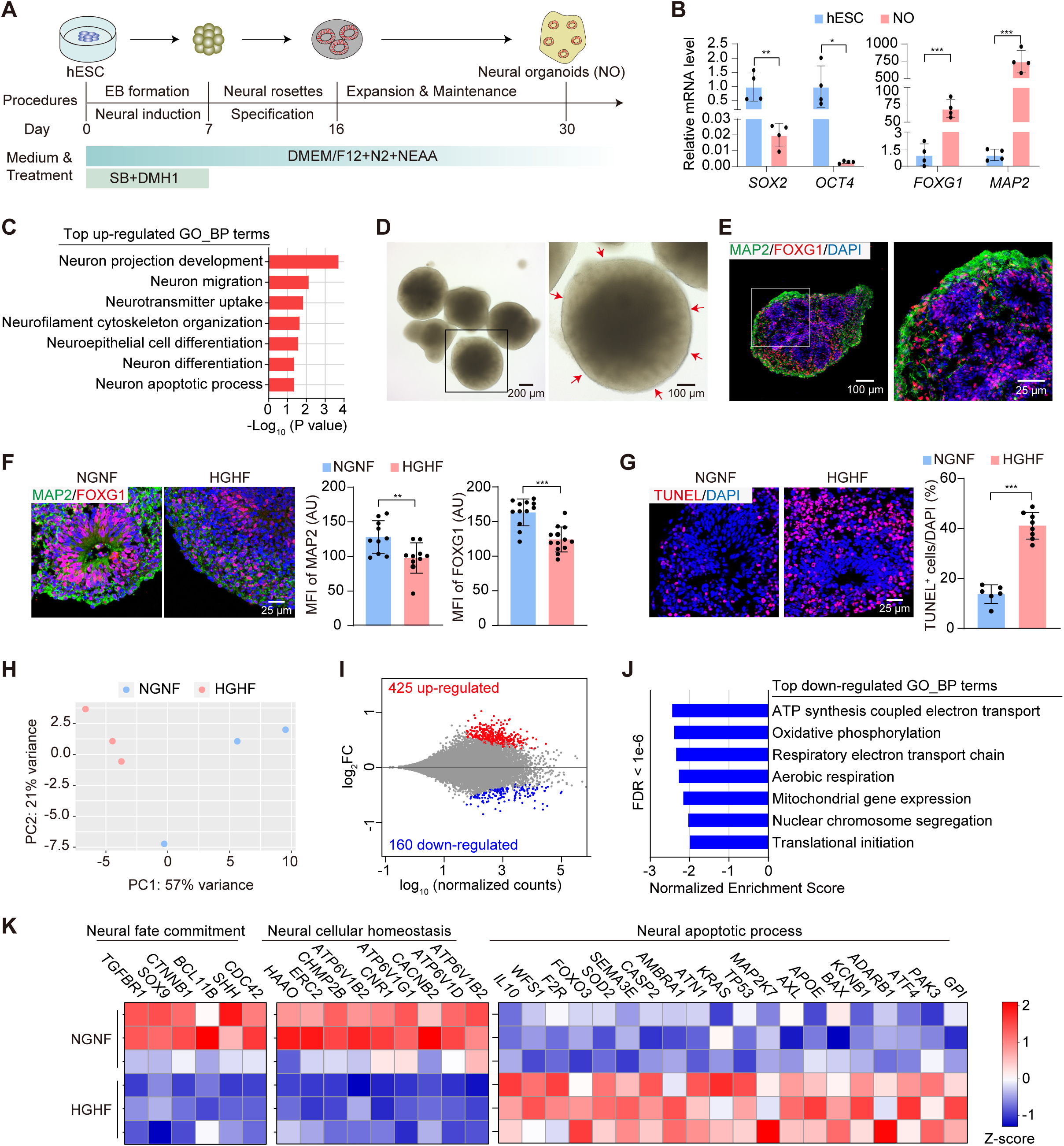
A glycolipotoxicity model of neural organoids. A) A flow chart of neural organoids (NOs) preparation. hESCs were cultured in suspension to form embryoid bodies (EBs) and treated with the TGF-β/Smad inhibitors SB431542 and BMP receptor inhibitor DMH-1 for 7 days to induce neural differentiation. Then the rosette structures were gradually formed, and NOs on day 30 were used for subsequent experiments. B) Real-time quantitative polymerase chain reaction (RT-qPCR) analysis of hESC markers (*SOX2, OCT4*) and neuronal markers (*FOXG1, MAP2*) in hESCs and NOs at day 30. C) Gene ontology-biological process (GOBP) analysis of top-500 highly expressed genes in NOs by Smart-seq. D) Representative bright-field images of NOs on day 30. E) Representative immunofluorescence images of the neural markers FOXG1 (red) and MAP2 (green) in NO sections. F-G) Immunofluorescence images and quantification of FOXG1/MAP2 (F) or TUNEL (G) staining in NOs treated with NGNF or HGHF medium for 2 days. MFI, mean fluorescence intensity. H-I) PCA plot (H) and MA plot (I) of Smart-seq analysis in NOs treated with NGNF or HGHF medium for 2 days. J) GOBP analysis of differentially down-regulated genes (FDR < 1e-6) in NOs treated with NGNF or HGHF medium for 2 days. K) Heatmap of gene expression involved in neural fate commitment, neural cellular homeostasis, and neural apoptotic process in Smart-seq data. In B, F and G, Data are plotted as mean ± SEM and analyzed by unpaired t test. *P < 0.05, **P < 0.01, ***P < 0.001.

NOs were next treated with high concentrations of glucose and fatty acid that were the same to the previous EHT treatment. Notably, the control NO medium involved 17.5 mM glucose and no palmitic acid, which was not identical to EHT control. HGHF profoundly decreased FOXG1 and MAP2 immunostaining signals (Figure 2F) and increased the proportion of TUNEL^+^ cells (Figure 2G). Smart-seq2 analysis revealed prominent separation by principal component analysis (Figure 2H) and 585 differentially expressed genes, 425 up-regulated and 160 down-regulated (Figure 2I, Supplementary Data 2). GO term analysis showed down-regulated ATP synthesis, oxidative phosphorylation and other oxidative respiration terms (Figure 2J). In addition, HGHF treatment down-regulated genes in neural fate commitment and neural cellular homeostasis, and up-regulated neuronal apoptosis (Figure 2K), which confirmed immunostaining analyses. These data support the successful establishment of an NO-based neural glucolipotoxicity model.

### 3.3 Exploration of an EHT-NO co-culture system

EHT and NO were routinely maintained using media with completely non-overlapping components (Figure 3A), partly due to empirical practice by the different laboratories. Thus, a nonnegligible task in building a co-culture system is to unify the cell culture medium. Based on existing knowledge about each component in the source media, three items in NO or EHT medium were combined to generate a generalized serum-free medium called general medium (GM) (Figure 3A).

**Figure 3.**
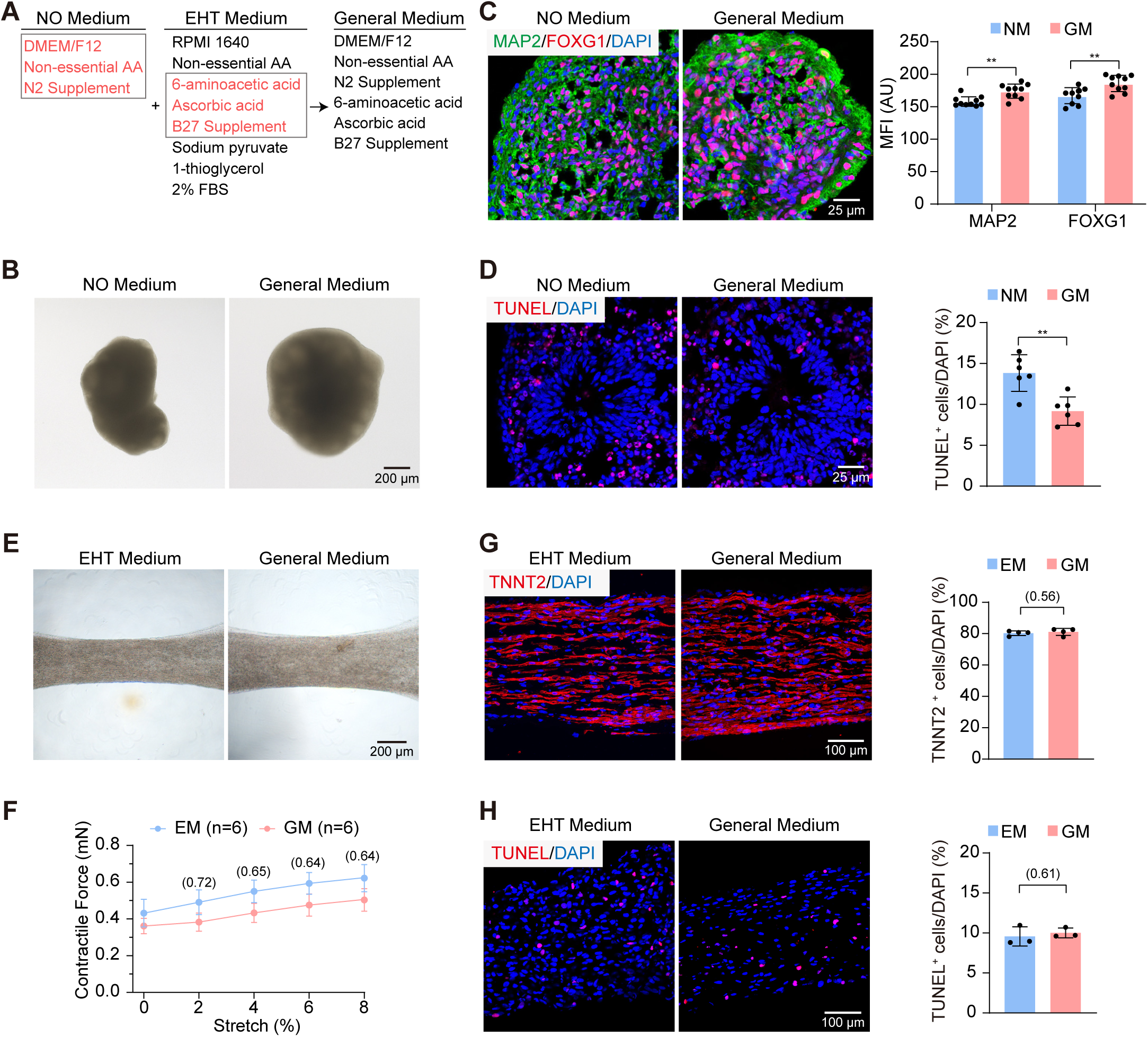
Development and validation of a general medium for EHTs-NOs co-culture. A) Components of neural medium (left), EHT medium (middle) and general medium for co-culture (right). Ingredients constituting the general medium are indicated in red. AA, amino acid. FBS, fetal bovine serum. B) Representative bright-field images of NOs cultured in neural medium or general medium for one week. C-D) Representative immunofluorescence images and quantification of FOXG1/MAP2 (C) or TUNEL (D) staining in NOs cultured in neural medium or general medium for one week. E) Representative bright-field images of EHTs cultured in the indicated medium for one week. F-G) Representative immunofluorescence images and quantification of TNNT2 (F) or TUNEL (G) staining in EHTs cultured in the indicated medium for one week. H) Quantitative statistics of contractile forces of electrically stimulated (3 HZ) EHTs cultured in the indicated medium for one week. Data are plotted as mean ± SEM. In C, D, G and H, Data are analyzed by unpaired t test. In F, Data are analyzed by two-way ANOVA with Tukey’s post hoc test. **P < 0.01.

As compared to the original NO medium (NM), the GM could maintain NOs for at least a week with no loss of morphology (Figure 3B). The levels of the neural markers FOXG1 and MAP2 were significantly increased (Figure 3C) while the proportion of apoptotic cells reduced (Figure 3D). Thus, GM outperformed the NO medium in NO maintenance. Next, the original EHT medium (EM) and GM were compared in culturing EHTs, which also revealed no changes in morphology (Figure 3E). EHTs exhibited similar contractile forces between EM and GM treatment (Figure 3F). The high proportion of TNNT2^+^ cardiomyocytes (Figure 3G) and the low level of TUNEL^+^ cells (Figure 3H) in EHTs could be maintained for more than a week by GM.

Based on the general medium, which contains 17.5 mM glucose and no palmitic acid, EHTs were next individually cultured or co-cultured with NOs for 48 h. In this condition, NOs did not alter the proportion of cardiomyocytes in EHTs (Figure 4A). The proportion of apoptotic cells was also unchanged (Figure 4B). The contractile forces showed no significant difference upon NO co-culture (Figure 4C-D). These results indicated that NOs had little effect on EHTs under the NGNF co-culture condition.

**Figure 4.**
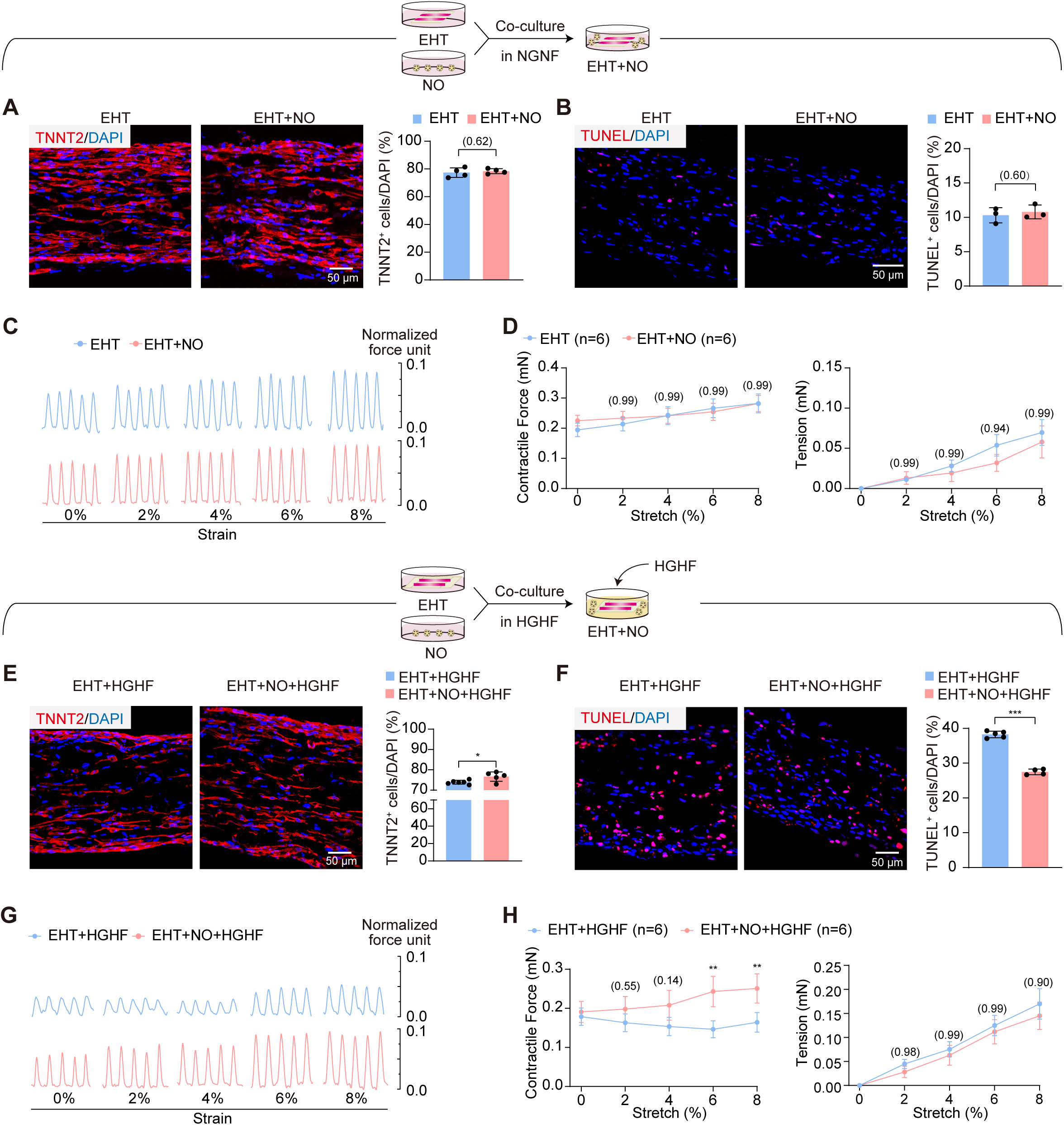
EHTs-NOs co-culture alleviates EHTs glucolipotoxicity in high glucose and high fatty acid (HGHF) treatment. A-B) Representative immunofluorescence images and quantification of TNNT2 (A) or TUNEL (B) staining of EHTs under NGNF culture for 24h. C-D) Contractile traces (C) and quantification of forces (D) of EHTs under NGNF culture for 24h. E-F) Representative immunofluorescence images and quantification of TNNT2 (E) or TUNEL (F) staining of EHTs under HGHF culture for 24h. G-H) Contractile traces (G) and quantification of forces (H) of EHTs under HGHF culture for 24h. Data are plotted as mean ± SEM and analyzed by unpaired t test (A, B, E, F) or two-way ANOVA with Tukey’s post hoc test (D, H). *P < 0.05, **P < 0.01, ***P < 0.001.

Then the high-glucose and high-fatty acid treatment was applied to this co-culture system. Interestingly, in comparison with HGHF-treated EHTs alone, co-culture with NOs allowed more TNNT2^+^ cardiomyocytes (Figure 4E) and less apoptotic cells (Figure 4F) to be detected in EHTs. The contractile forces of EHTs were not restored at the basal condition but become improved when strains were applied (Figure 4G-H). However, there was no significant difference in diastolic tension (Figure 4G-H). These results indicated that NOs could ameliorate myocardial glucolipotoxicity.

### 3.4 Validation of EHT-NO co-culture using human EHTs

The above studies were based on rat EHTs. To test if the effects were conserved in human cells, cardiac cells were next derived from hESCs to assemble into human EHTs using an adapted protocol^23^ (Figure 5A). In this protocol, TNNT2^+^ cardiomyocytes composited ∼70% total cell population (Figure 5B). Stem cell markers *SOX2* and *OCT4* were significantly decreased while cardiac markers *TNNT2* and *ACTN2* were significantly increased during hESC differentiation (Figure 5C). Human EHTs also endowed cardiomyocytes the elongated morphology (Figure 5D) and strain-induced contractility enhancement (Figure 5E-F).

**Figure 5.**
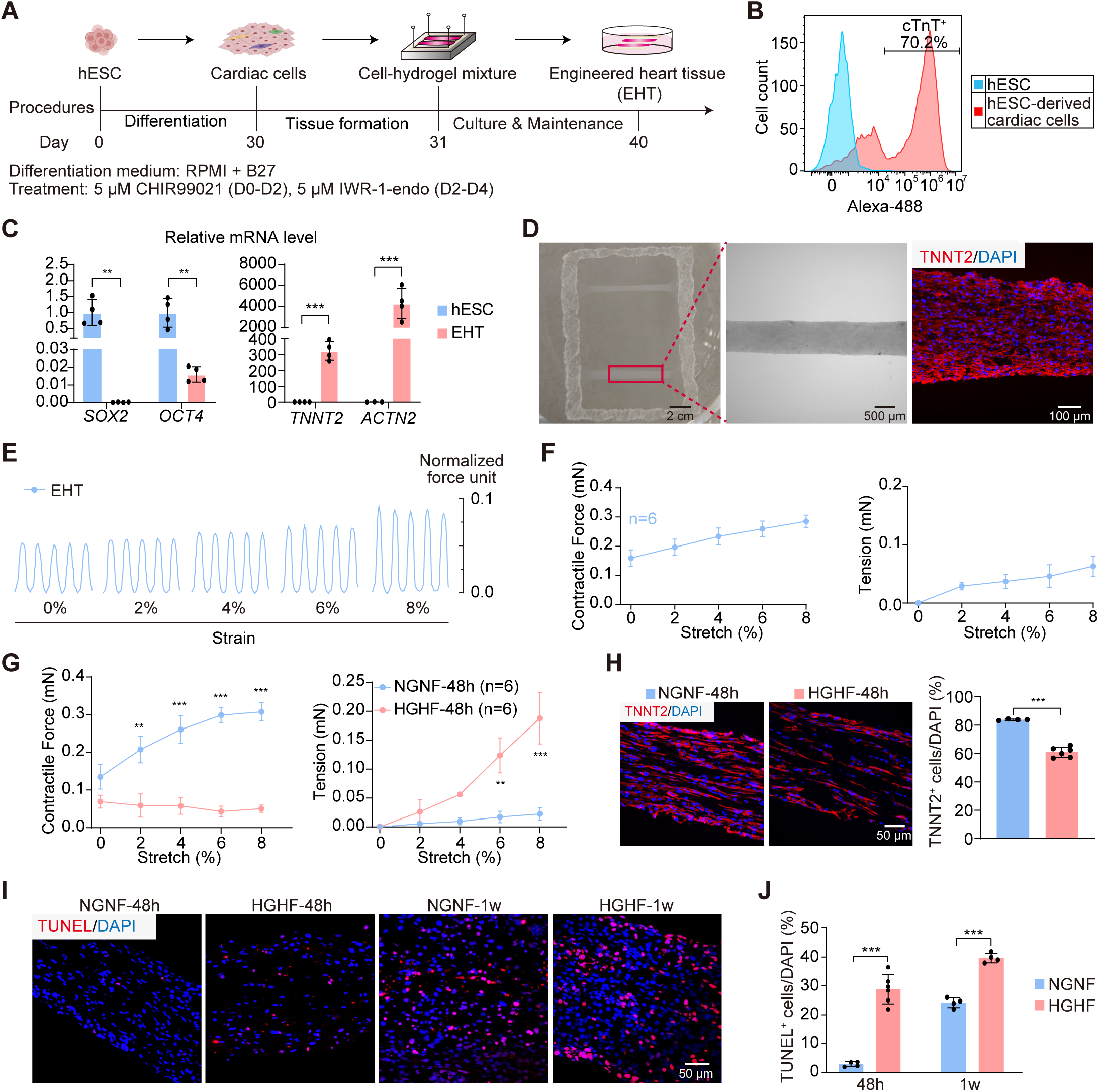
Verification of the EHTs-NOs co-culture effect using human embryonic stem cells (hESCs)-derived EHTs. A) A flow chart of hESCs-derived EHTs preparation. hESCs were differentiated into cardiac cells by sequential treatment of ChIR99021 and IWR-1-endo. The beating cells were then matured as a monolayer until day 30 before they were dissociated and casted into EHTs similar to the procedure in figure 1A. These human EHTs were cultured for 10 days before subsequent experiments. B) The ratio of cardiomyocytes versus non-cardiomyocytes was measured by flow cytometry. C) RT-qPCR quantitative analysis of stem cell markers (*SOX2, OCT4*) and cardiac markers (*TNNT2, ACTN2*) in hESCs and EHTs. D) Representative bright-field images of human EHT and immunofluorescence staining of TNNT2 (red) and nuclear DAPI (blue). E-F) Contractile traces and force quantification of hEHTs under electrical stimulation (1.5 HZ) during progressive stretching. G) Quantification of the contractile force and passive tension of human EHTs under NGNF or HGHF treatment for 48 h. H) Representative images and quantification of TNNT2 in human EHTs under NGNF or HGHF treatment for 48h. I-J) Representative images (I) and quantification (J) of TUNEL staining in human EHTs under NGNF or HGHF treatment for 48 h or one week (1w). Data were plotted as mean ± SEM and analyzed by unpaired t test (B, C, H, J) or two-way ANOVA with Tukey’s post hoc test (G). **P < 0.01, ***P < 0.001.

Human EHTs generally exerted weaker forces than rat EHTs (Figure 1F versus Figure 5F), indicating differences in cardiomyocyte maturity or quality^40^. Human EHTs were next treated with HGHF general medium for 48 h, which completely abolished contractility while increased tissue stiffness (Figure 5G). HGHF treatment caused decreased TNNT2^+^ cardiomyocytes and increased TUNEL^+^ cells in human EHTs at 48 h, which was aggravated as the treatment was extended to 1 week (Figure 5H-J). Since 48 h treatment was sufficient to trigger a severe phenotype while extended 1-week culture also increased cell death in control (Figure 5I-J), the following studies were focused on 48 h of HGHF treatment.

Next, human EHTs were subjected to RNA-seq analysis (Supplementary Data 3) and compared to rat EHTs in the HGHF treatment. Homologous genes were identified in the two species via BioMart community portal^41^ and an overlap analysis of differentially expressed homologs revealed more than 60% distinction (Figure 6A). A representative example included the *MYH6-7* isoform switching, which was known to demonstrate opposite changes in cardiac diseases in human versus rodents (Figure 6B)^40^. Unlike single-gene analyses, however, GSEA and pathway analyses demonstrated at least 70% overlap between human and rat (Figure 6C), which included typical glucolipotoxicity pathways including increased cytokine pathways and apoptosis, and decreased oxidative respiration and muscle contraction (Figure 6D).

**Figure 6.**
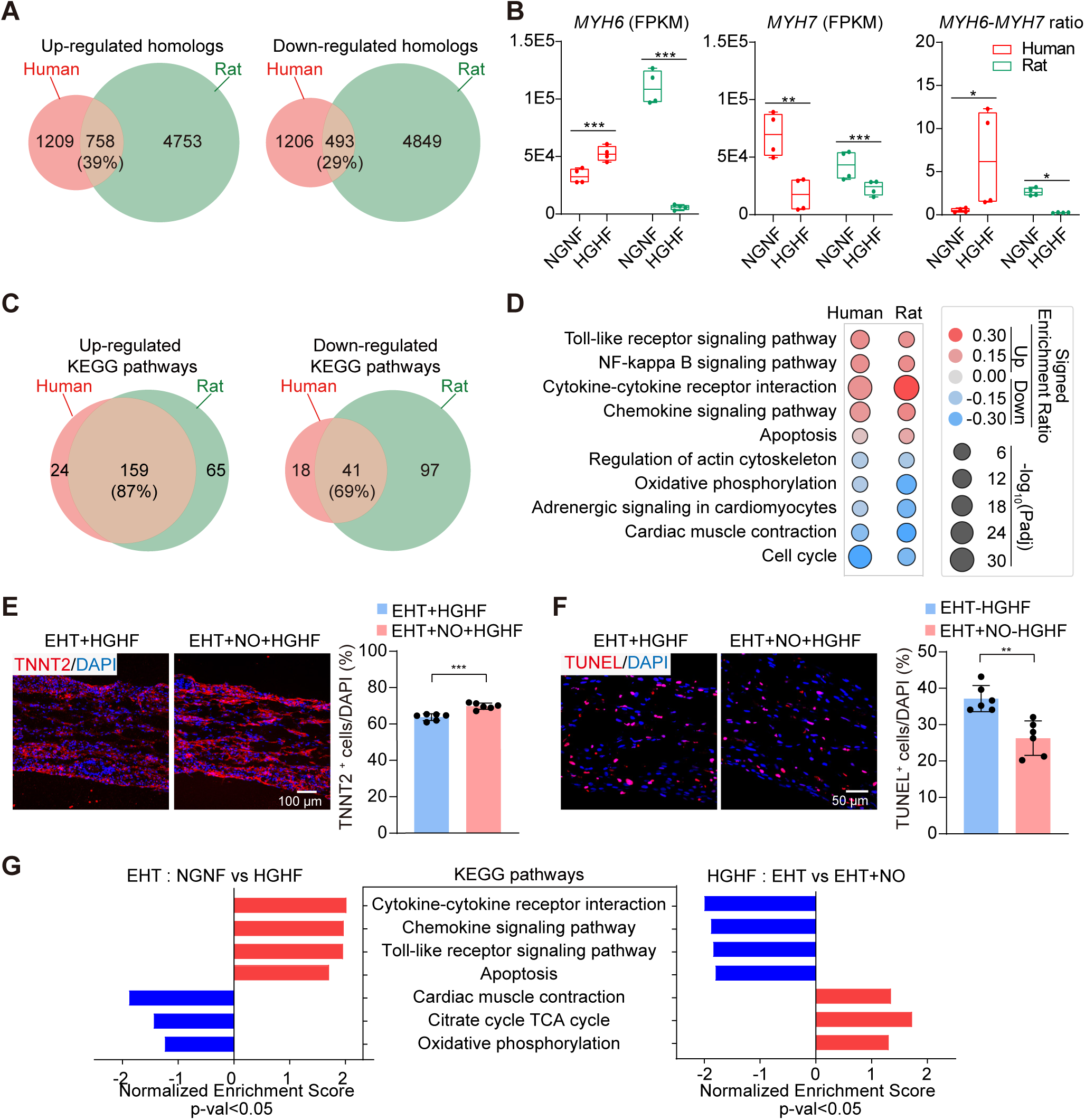
Comparative EHT RNA-seq analysis between human and rat. A) Venn diagram of differentially expressed homologous genes (Padj < 0.05) in human versus rat EHTs upon HGHF treatment. Homologous genes were identified using the BioMart database. B) Species-specific expression changes of *MYH6, MYH7* and their ratios. Statistical analysis via DESeq2 or Mann-Whitney U test. C) Venn diagram of significantly altered KEGG pathways (Padj < 0.05) in human versus rat EHTs upon HGHF treatment. D) Representative KEGG pathways that were correlatively altered in human versus rat EHTs upon HGHF treatment. KEGG analysis was performed via KOBAS-i. E-F) Confocal images and quantification of TNNT2 (E) or TUNEL (F) in human EHTs in individual culture versus NO co-culture under the HGHF treatment for 48 h. G) Smart-seq2 analysis and KEGG pathway enrichment in comparison between NGNF and HGHF or between individual culture and NO co-culture. Up-regulated gene sets in red and down-regulated in blue. Data were plotted as mean ± SEM and analyzed by unpaired t test (E, F). *P<0.05, **P < 0.01, ***P < 0.001.

Next, human EHTs were co-cultured with NOs of the same hESC origin and treated with HGHF general medium. The proportion of TNNT2^+^ cardiomyocytes in EHTs in the co-culture group was significantly higher than that in the EHTs that were cultured alone (Figure 6E). The proportion of TUNEL^+^ cells in EHTs was also significantly reduced by NO co-culture (Figure 6F). To further confirm this effect, RNA-seq analysis of EHTs in EHT-NO co-culture demonstrated a clear reversal of glucolipotoxicity gene sets (Figure 6G , Supplementary Data 4). Therefore, the protective effect of NOs on EHTs is independent of the EHT species.

### 3.5 Mechanistic investigation of NO-to-EHT signaling in co-culture

Because NOs and EHTs were not physically contacted in co-culture, RNA-seq of NOs was revisited to identify potential secretory factors that mediated this communication. An array of neurotrophic factors, extracellular matrix components and inflammatory cytokines were differentially regulated between NOs in NGNF versus HGHF conditions (Supplementary Figure 2). The differentially regulated genes in NOs were next ranked by their expression levels. Among the top 10 upregulated NO genes, *VCAN* was the only gene coding a secretory protein (Figure 7A). Published single-cell RNA-seq data ^26^ showed broad expression of *VCAN* in most cell populations in NOs (Supplementary Figure 3), among which intermediate progenitor cells expressed the highest level of *VCAN* as expected, since they were considered as stromal cell types secreting extracellular matrix proteins^42^.

**Figure 7.**
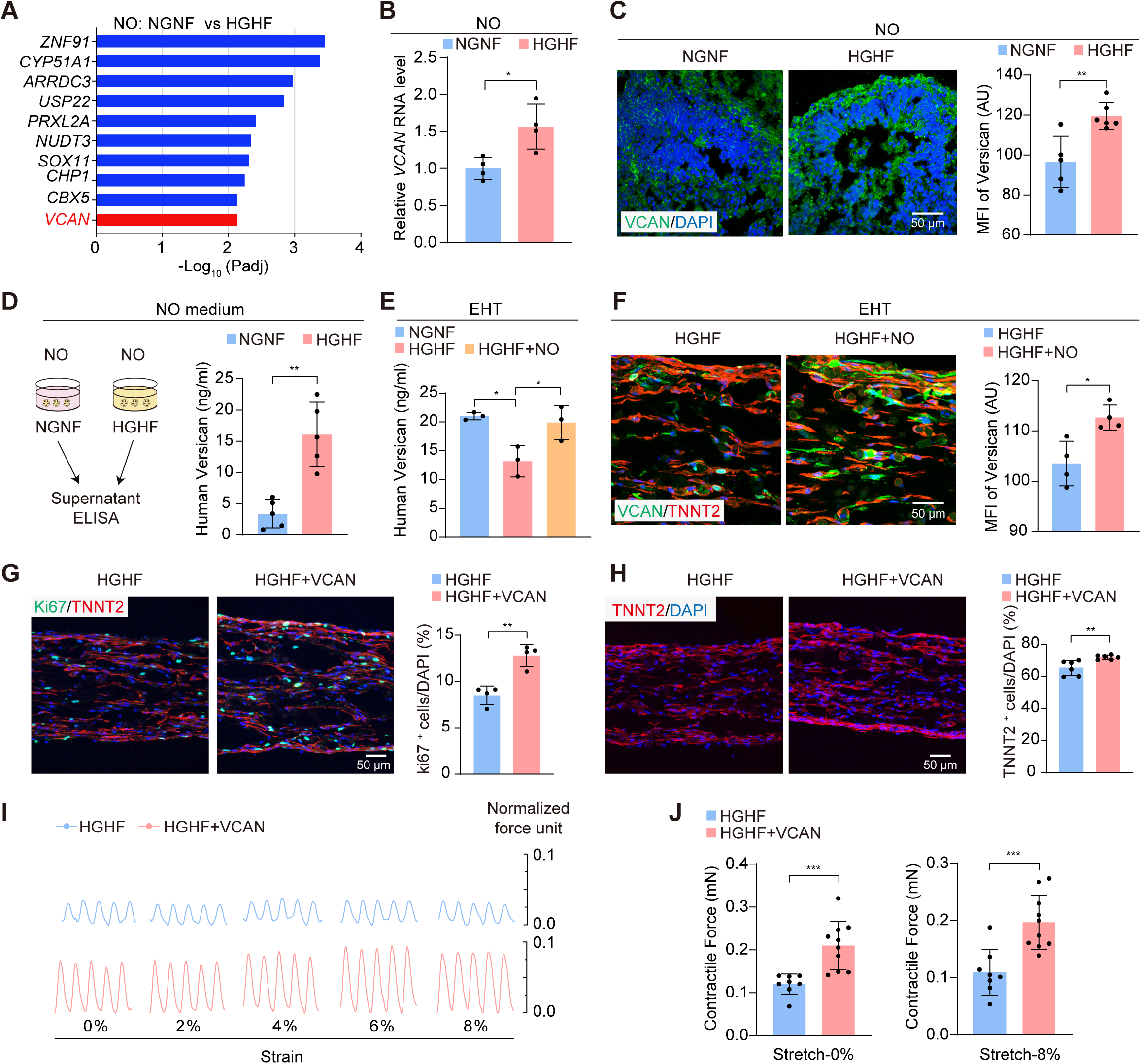
NOs transfer versican into EHTs to ameliorate myocardial glucolipotoxicity in co-culture. A) Top-10 highly expressed genes that were significantly up-regulated in NOs treated with HGHF. Screening criteria: Padj < 0.05, log2FC > 0 and Basemean > 3500. The only one encoding a secreted protein was *VCAN* (red). B) RT-qPCR of NOs showing transcriptional upregulation of *VCAN* in NOs treated with HGHF for 48 h. C) Confocal images and quantification of VCAN (green) in NGNF and HGHF-treated NOs. D) ELISA-based detection of supernatant versican in NGNF- and HGHF-treated NO groups. E) Quantitative analysis of versican protein levels in EHT lysates measured by ELISA. F) Representative immunostaining images of VCAN (green) and TNNT2 (red) and quantitative analysis of VCAN mean fluorescence intensity (MFI) in EHT individual and co-culture groups. G-H) Representative immunofluorescence images and quantification of cell proliferation marker ki67 (green) and TNNT2 (red) in EHT and EHT + 10 ng/ml versican groups under the HGHF treatment for 48 h. I-J) Representative contractile traces (I) and quantification (J) of EHT contractility in the indicated groups. Data are plotted as mean ± SEM and analyzed by unpaired t test (B, C, D, F, G, H, J) or one-way ANOVA with Tukey’s post hoc test (E). *P < 0.05, **P < 0.01, ***P < 0.001.

The upregulation of *VCAN* in NOs by HGHF were next validated by quantitative PCR (Figure 7B) and immunofluorescence (Figure 7C). Enzyme-linked immunosorbent assay (ELISA) detected a 3-fold increase of versican in NO culture medium with HGHF (Figure 7D). By contrary, the expression of versican was decreased by HGHF treatment in the EHTs, while NO increased versican in EHTs in co-culture (Figure 7E). Immunofluorescence confirmed the elevated versican in EHTs upon co-culture with NOs (Figure 7F). These data indicated that NOs transferred versican to EHTs in the co-culture under HGHF treatment.

Versican was reported to promote cell proliferation and myocardial regeneration in a murine myocardial infarction model^43^. To determine if a similar effect was applied to EHTs, 10 ng/ml versican recombinant protein was applied to EHTs and a significant increase of ki67^+^ proliferative cells was observed (Figure 7G). The proportion of TNNT2^+^ cardiomyocytes was also increased by versican (Figure 7H), which was associated with partially restored contractile forces (Figure 7I-J). Versican also alleviated dysregulation in calcium transients in cardiomyocytes under HGHF treatment (Supplementary Figure 4, Supplementary Video 2-4). Thus, versican could mediate the NO-based amelioration of myocardial glucolipotoxicity in EHTs.

### 3.6 Versican improves cardiac function in diabetic cardiomyopathy in mice

Subsequently, a diabetic cardiomyopathy (DbCM) mouse model was used to test the effect of versican on the heart in vivo. We first established DbCM mice via a high-fat-diet (HFD)-Streptozotocin (STZ) treatment protocol as previously described^44^ (Figure 8A). Increased body weight and impaired tolerance to glucose and insulin validated metabolic dysfunction in these mice (Supplementary Figure 5A-C). Elevated blood glucose, serum free fatty acid and total cholesterol was also confirmed (Figure 8B-D).

**Figure 8.**
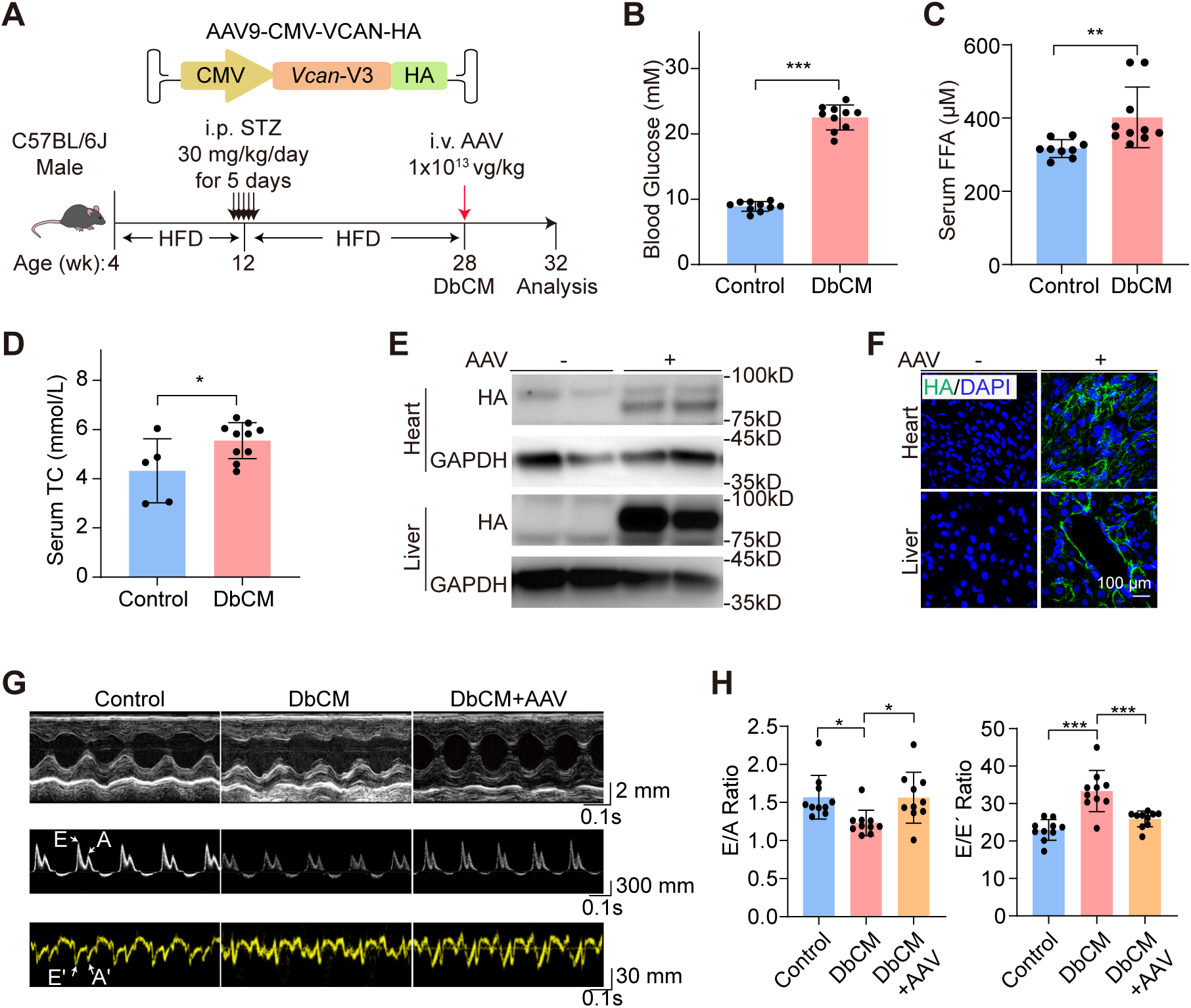
Versican improves cardiac function in diabetic cardiomyopathy in mice. A) Schematic diagram of the generation of DbCM mice and AAV treatment. B-D) Measurements of fasting blood glucose (B), serum free fatty acids (FFA)(C) and total cholesterol (TC)(D). E) Western blot of AAV-delivered VCAN expression by HA tag. F) Immunofluorescence images of VCAN-HA. G) Representative parasternal short-axis echocardiography (top), Pulsed-Wave Doppler (middle) and Tissue Doppler images (bottom). H) Quantitative analysis of the ratio of flow Doppler E wave amplitude to A wave amplitude (E/A) and ratio of flow Doppler E wave amplitude to tissue Doppler E′ wave amplitude (E/E′). Data are plotted as mean ± SEM and analyzed by unpaired t test (B, C, D) or one-way ANOVA with Tukey’s post hoc test (H). *P < 0.05, **P < 0.01, ***P < 0.001.

Adeno-associated virus 9 (AAV9) was generated to globally express VCAN-V3^45^, the only natural versican isoform that could fit into the limited payload of AAV (Figure 8A). Western blot validated AAV-delivered VCAN expression in the heart and liver (Figure 8E, Supplementary Figure 6). Immunofluorescence imaging confirmed the extracellular fibrous structures of AAV-expressed VCAN as an ECM protein (Figure 8F). Next, 1×10^13^ vg/kg AAV was intravenously injected into 28-week-old DbCM mice. After 4 weeks, the mice were subjected to echocardiography, which detected restored ejection fraction, fractional shortening (Supplementary Figure 5D-E), E/A ratio and E/E’ ratio (Figure 8G-H). AAV did not alter blood free fatty acid level (Supplementary Figure 5F), suggesting little indirect impact on lipid metabolism. These data confirmed the organoid co-culture study and indicated the protective role of versican in DbCM.

## 4. Discussion

Metabolic disorders could deposit glucolipotoxicity at the systemic level^46–48^, but how distal inter-organ communications contribute to pathogenesis remains poorly understood. Animal models are complicated by the presence of all the organs, which is difficult to dissect communications between individual organs, thus a simplified in vitro system with designated tissues and organs are necessary. This study utilized EHTs and NOs as an example to demonstrate a workflow to build and study inter-organ communications and provided proof-of-concept evidence about the presence of material transfer between organoids and engineered tissues that could influence pathological phenotypes related to glucolipotoxicity.

In the last decade, tissue engineering and organoid technologies are rapidly evolving, but co-culture or assembloid studies have been limited to organs and tissues of close anatomical or developmental relationship^15, 18, 49^. The combination of more distal organoids was challenging partly due to the distinct protocols to build and maintain each organoid. For example, an immediate challenge to co-culture cardiac and neural organoids was to generalize the co-culture media. In this study, the original NO medium lacked 6-aminoacetic acid, which was an essential component to build EHT^23^, while the original EHT medium not only contained serum but also lacked the N2 supplement for NO maintenance. To solve this problem, we carefully evaluated and selected essential components to establish a unified, chemically defined medium with comparable or even better capacity to maintain EHTs and NOs together.

Neuronal regulation of the heart is thought to occur through the effects of innervation and neurotransmitters^50–52^, thus scientists naturally believe that NOs and EHTs should physically contact to build a microphysiological system with neural innervation on muscle cells. However, in such a potential model, it would be challenging to dissect between the innervated and secretory effects. Therefore, in this study, we purposely positioned NOs and EHTs in co-culture without physical contact, which indeed justified the presence of non-negligible secretory effects from NOs to EHTs. These results are crucial reference in future studies that try to build biophysical connection between NOs and EHTs.

Versican is an important chondroitin sulfate proteoglycan found in the extracellular matrix. Versican is canonically believed to be expressed by fibroblasts and play essential roles for cardiac development and repair^43, 53–55^. Interestingly, this study showed that EHTs did not increase versican by themselves but received versican from NOs in co-culture. In addition, versican treatment to EHTs is sufficient to recapitulate NO-medicated protective effect against glucolipotoxicity. These data suggested that myocardium could potentially receive ECM components from extracardiac sources to modify pathogenic outcomes. These findings also provide evidence justifying the validity to acquire information with therapeutic potential in an organoid co-culture system.

Despite these interesting findings, several open questions remain to be answered in future studies. First of all, the cell source and molecular mechanism of glucolipotoxicity-induced versican release from NOs remain unclear. Single-cell characterization of NOs is necessary to solve this problem. Secondly, whether the NO-versican-EHT axis could be validated in animal models is also unclear. In the more complicated in vivo system, versican as a broadly expressed protein could potentially be transferred to the heart from other tissues rather than nerves. Above all, a versican-based therapeutic approach needs to be tested in animal models to examine if versican is a valid therapeutic target for diabetic cardiomyopathy.

## 5. Conclusion

A co-culture system was established to maintain and study NO-EHT communications in a dish. The glucolipotoxicity of EHTs could be alleviated by co-culture with NOs partly via versican transfer from NOs to EHTs. Versican overexpression could alleviate cardiac glucolipotoxicity in vivo.

## Author contributions

Y Guo and J Liu conceptualized and supervised the study. B Bai conducted and analyzed the NO study. J Li conducted EHT experiments. Z Wang conducted hESC differentiation. J He conducted rat cardiac cell isolation. Y Zhang, X Han, W Zhu and Y Liu generated NOs and helped with NO experiments. Y Qi, Z Wan, L Cai and D Zhang helped to build the EHT and glucolipotoxicity methodologies. Z Wang and R Wang performed calcium transient analysis. K Wang, J Zhang, W Huang, R Xu and M Sun provide advice and guidance on organoid co-culture and microphysiological system studies. Y Yang and D Zhao performed bioinformatic analysis. Y Guo, B Bai and J Li wrote and revised the manuscript.

## Acknowledgements

We thank Geekgene for Smart-seq analysis and Cellapy for technical assistance in hESC differentiation.

## Data availability statement

RNA sequencing data have been deposited in National Genomics Data Center (GSA-Human: HRA009640) that are publicly accessible at https://ngdc.cncb.ac.cn/gsa-human. AAV plasmid is available at Addgene or WeKwikGene. Other data that support the findings of this study are available on request via.

## Ethical statement

This study does not involve human patients. All animal experimental procedures were approved by the Institutional Animal Care and Use Committee of Peking University (approval number DLASBD0203).

