## Supplementary material for "Neural organoids protect engineered heart tissues from glucolipotoxicity by transferring versican in a co-culture system": Figure S1

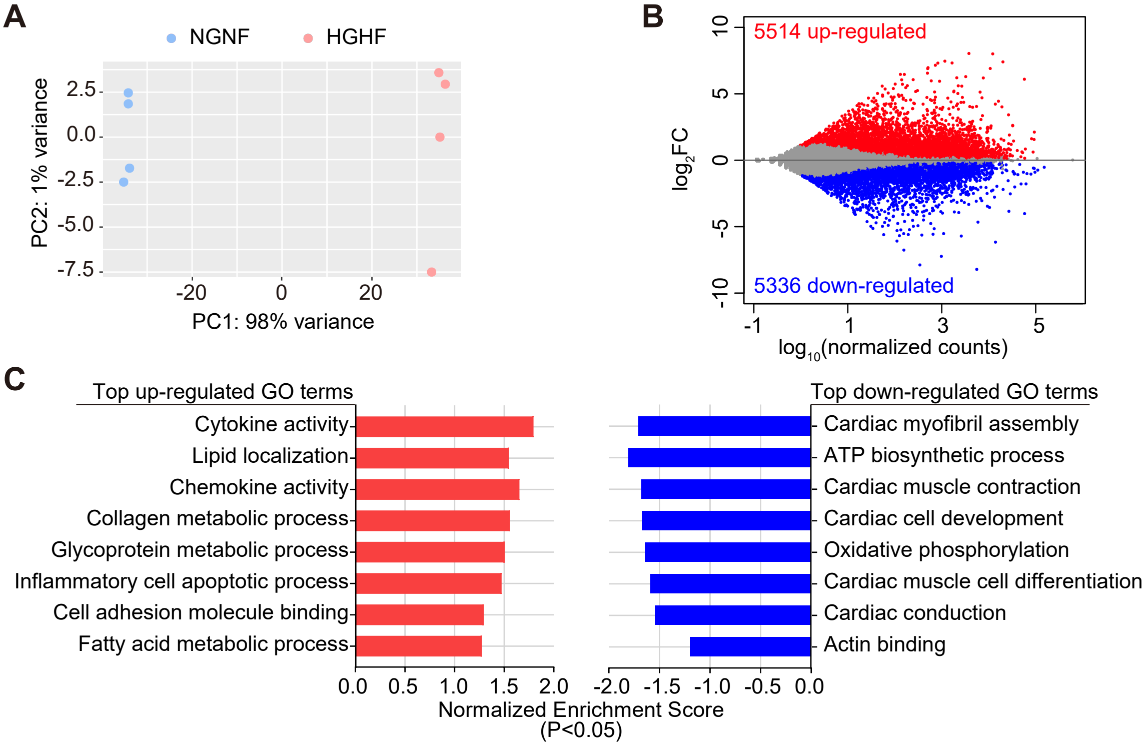


**Figure S1. RNA-seq analysis of glucolipotoxicity in rat EHTs.** A) PCA plot showing segregation of RNA-seq data. Rat EHTs were treated with NGNF or HGHF medium for 2 days. B) MA plot showing differentially expressed genes (Padj<0.05). C) GSEA analysis of differentially regulated GO terms in rat EHTs treated with NGNF or HGHF medium for 2 days.
