## Supplementary material for "Neural organoids protect engineered heart tissues from glucolipotoxicity by transferring versican in a co-culture system": Figure S2

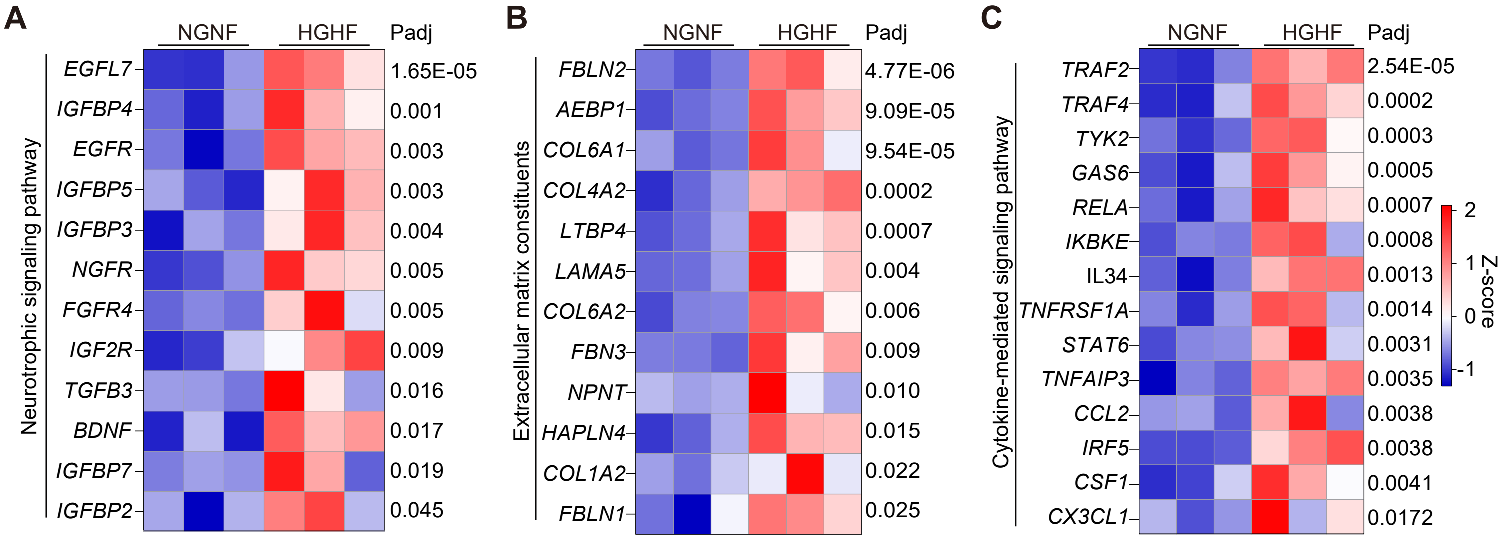


**Figure S2. Heatmap of Smart-seq analysis of NOs treated with NGNF or HGHF medium for 2 days.** A) Neurotrophic signaling pathway. B) Extracellular matrix constituents. C) Cytokine signaling pathways.
