## Supplementary material for "Neural organoids protect engineered heart tissues from glucolipotoxicity by transferring versican in a co-culture system": Figure S3

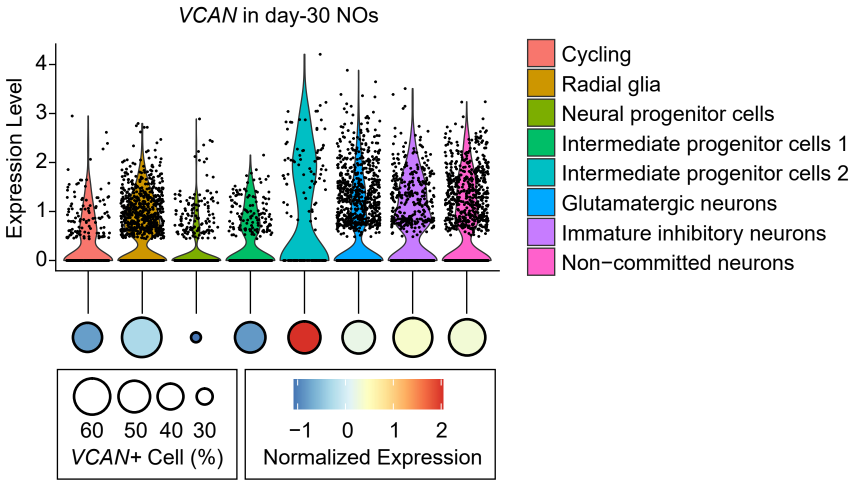


**Figure S3. Single-cell RNA-seq analysis of *VCAN* in day-30 NOs.** The violin plot and dot plot shows the expression levels of *VCAN* gene in different cell clusters. The bubble plot shows the proportion of *VCAN*-positive cells and the the expression levels of *VCAN* gene .
