## Supplementary material for "Neural organoids protect engineered heart tissues from glucolipotoxicity by transferring versican in a co-culture system": Figure S4

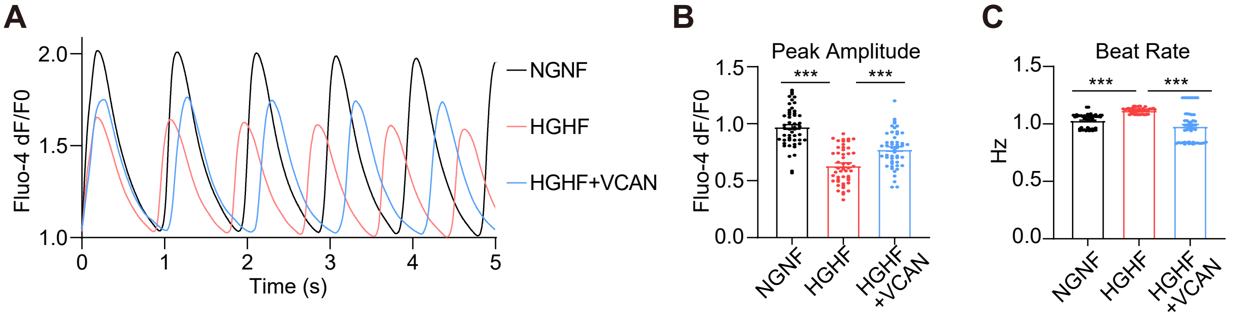


**Figure S4. The effect of versican on cardiomyocyte calcium transient.** A) Average $Ca^{2+}$ transeint signals of neonatal rat ventricular cardiomyocytes under autonomous beating. B) Analysis of $Ca^{2+}$ transeint peak amplitudes of NRVMs. C) Cardiomyocyte beat rate analysis. Data are plotted as mean ± SEM and analyzed by Mann-Whitney test (B, C), n = 50 cells per group, ***P<0.001.
