## Supplementary material for "Neural organoids protect engineered heart tissues from glucolipotoxicity by transferring versican in a co-culture system": Figure S5

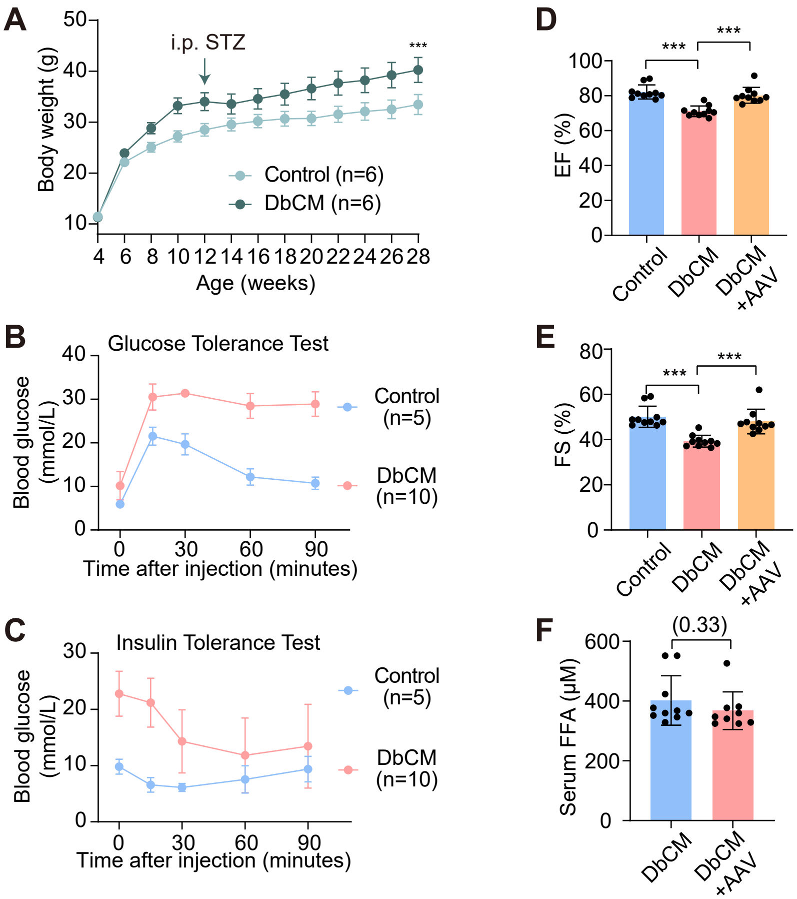


**Figure S5. Basic characterization of DbCM mice.** A) Body weight changes of mice in the control and DbCM group. B-C) Changes in blood glucose metabolism was detected by intraperitoneal glucose tolerance test (B) and intraperitoneal insulin tolerance test (C). D-E) Echocardiogram analysis of ejection fraction (EF) (D) and fractional shortening (FS) (E). F) Quantifcation of serum free fatty acids (FFA) concentration in DbCM and DbCM+AAV groups. Data are plotted as mean ± SEM and analyzed by two-way ANOVA with Tukey's post hoc test (A), one-way ANOVA with Tukey's post hoc test (D, E) or unpaired t test (F). *P < 0.05; **P < 0.01; ***P < 0.001.
