## Supplementary figures and images for "Neural organoids protect engineered heart tissues from glucolipotoxicity by transferring versican in a co-culture system"

### Figure S6

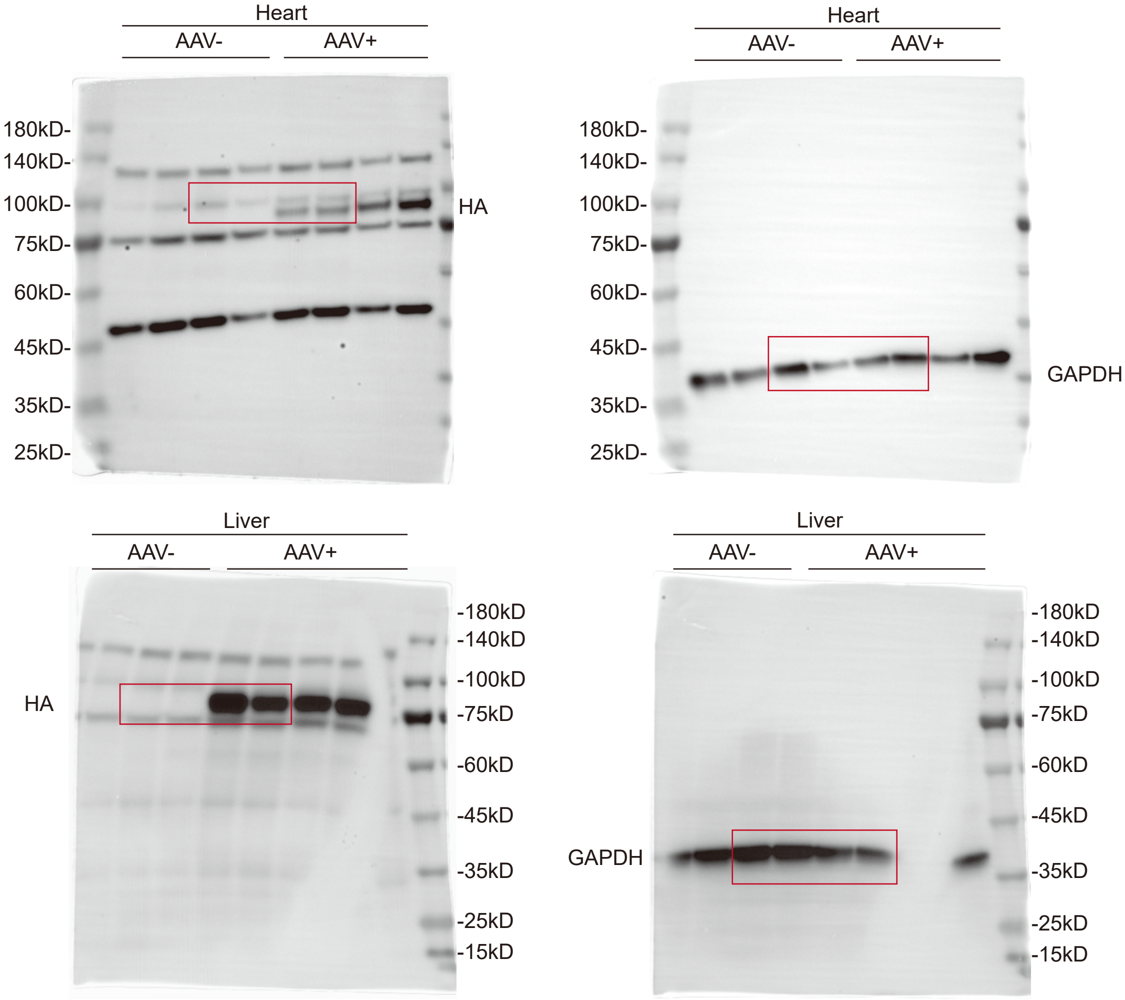


**Figure S6. Uncropped Western blot images for Figure 8E.**
